# CSF circulation and dispersion yield rapid clearance from intracranial compartments

**DOI:** 10.1101/2022.05.02.490257

**Authors:** Martin Hornkjøl, Lars Magnus Valnes, Geir Ringstad, Marie E. Rognes, Per-Kristian Eide, Kent-André Mardal, Vegard Vinje

## Abstract

In this paper we used a computational model to estimate the clearance of tracer driven by circulation of cerebrospinal fluid (CSF) produced in the choroid plexus (CP) located within the lateral ventricles. CSF was assumed to exit the subarachnoid space (SAS) via different outflow routes such as the parasagittal dura, cribriform plate and/or meningeal lymphatics. We also modelled a reverse case where fluid was produced within the spinal canal and absorbed in the CP in line with observation on certain iNPH patients. No directional interstitial fluid flow was assumed within the brain parenchyma. Tracers were injected into the foramen magnum. The models demonstrate that convection in the SAS yield rapid clearance from both the SAS and the brain interstitial fluid (ISF) and can speed up intracranial clearance from years, as would be the case for purely diffusive flow, to days.

## 1 Introduction

Cerebrospinal fluid (CSF) flow plays a fundamental role in the clearance of solutes from intracranial compartments^1, 2^. Current views postulate that CSF is primarly produced in the choroid plexus^1, 3^, and flows through the ventricular system^4–6^ and along the subarachnoid space (SAS)^7–9^. From there, CSF drains towards the venous system via arachnoid granulations^10^, towards lymph nodes via e.g. perineural routes across the cribriform plate^2, 7, 11^ or the meningeal lymphatics^12^, or flows through the brain parenchyma itself via glymphatic (perivascular) pathways^13^. The relative importance of these pathways, their interplay, and role(s) in physiological as well as pathological solute transport remain unresolved^1, 2, 7, 8, 10, 12, 14^.

Importantly, CSF circulation characteristics change under physiological transitions, in neurological disorders and with neurodegenerative disease. In patients diagnosed with idiopathic normal pressure hydrocephalus (iNPH), MR-imaging reveals altered solute influx and clearance rates^15^. In both Alzheimer’s and iNPH patients, CSF dynamics in the SAS is altered^15, 16^, and CSF production within the choroid plexus may be reduced in iNPH^6^. On the other hand, changes in glymphatic function may be associated with several types of dementia^17^. In Alzheimer’s disease, alterations in arterial pulsatility^18^, AQP4 function^19^ and sleep disturbances^20^ has been proposed as causes of glymphatic impairment. Lastly, glymphatic transport has been reported to increase during sleep^21, 22^.

A key question is to what extent the CSF circulation induced by CSF production, vascular pulsatility and CSF efflux contributes to transport of solutes (both influx and outflux) in the SAS and brain parenchyma. While intraparenchymal transport and glymphatics have received substantial attention over the last decade^1, 13, 14, 21–29^, the clearance interplay between different regions within the intracranial compartment is less understood. To illustrate, while Xie et al^21^ suggest that the sleep-wake cycle regulates the efficiency of glymphatic solute clearance via changes in the interstitial space volume, the findings of Ma et al^7^ offer an alternative interpretation in which increased CSF outflux during wakefulness effectively limits the availability of solutes at the surface and within parenchymal perivascular spaces (PVSs). As the intracranial CSF volume is only 10–30% that of the brain^30, 31^, rapid clearance of substances from the SAS is crucial to sustain diffusive transport from the brain parenchyma to the SAS.

Crucially, CSF flow velocities in the SAS, including in surface PVSs, are substantial. Pulsatile CSF velocities of at least 10–40 *μ*m/s can be inferred from experimental measurements of microsphere movement in rodents^8, 9^. Furthermore, the resulting dispersion effects may dominate diffusion by a factor of 10^4^ for transport of smaller molecules such as the MRI contrast molecule Gadoteridol^29^. In humans, CSF flow in the SAS varies significantly between patients and diseases^32^, with velocities at the foramen magnum induced by pulsatile flow on the order of 5 cm/s^33^. Interestingly, CSF bulk flow at a magnitude of *μ*m/s can be induced in the ventricular system and surface PVSs by relatively small intracranial pressure gradients (< 1–2 mmHg/m)^34^.

In this study, using biophysics-based finite element computational models created from T1- and T2-weighted MR images^15, 35^, we study CSF flow in the ventricular system and SAS and solute transport in these CSF-filled spaces and brain parenchyma. We first simulate flow patterns and magnitude induced by a production of 0.5L CSF per day^36^ in the choroid plexus and different CSF efflux pathways: across the parasagittal dura, across the cribriform plate, and into meningeal lymphatics, as well as reversed flow scenarios. We next simulate solute transport in the SAS and brain parenchyma resulting from an intrathecal injection of Gadobutrol. Our findings indicate that CSF flow in the SAS is a major player in brain clearance. However, no single outflow pathway alone is able to explain in-vivo observations of brain-wide distribution of tracer combined with fast clearance from the SAS, and we thus propose that a combination of different outflow routes seem more likely.

## 2 Methods

In this computational study, we quantify and characterize CSF flow patterns and molecular transport in the SAS and parenchyma induced by different clearance pathways. We also consider a choroid plexus-based production of 0.5 L/day of CSF and efflux across the 1) parasagittal dura^37^, 2) the cribriform plate^11^, and 3) meningeal lymphatics^12^. We consider a scenario with retrograde flow in the aqueduct^4^ by assuming that 0.5 L/day CSF production occurs within the spinal cord and as such that there is a influx through the foramen magnum, combined with an efflux route in the choroid plexus. An illustration of a slice of the computational domain is given in Figure 1.

**Figure 1.**
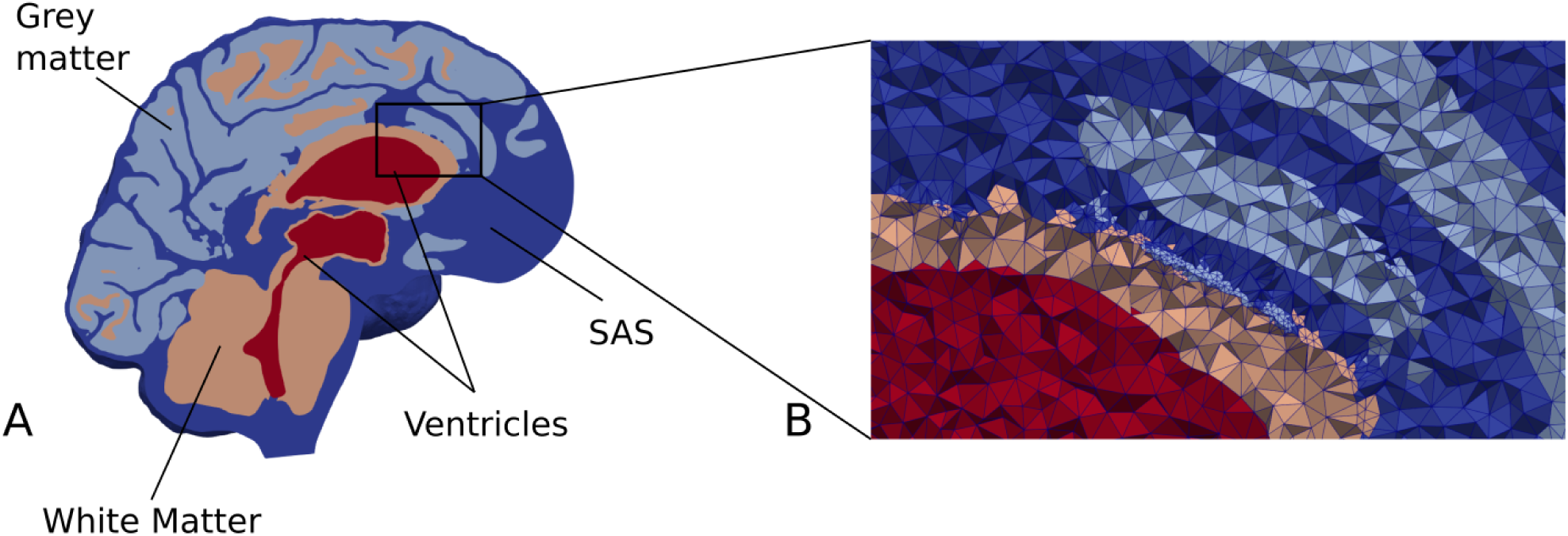
**(A)** A cross section of our brain mesh showing the SAS (dark blue), white matter (orange), gray matter (light blue), ventricles (red), **(B)** shows a zoom in on a part of the mesh with the edges of the mesh triangles. Note that for visualization purposes, the resolution shown here is coarser than the resolution used in the numerical simulations.

### 2.1 Patient data and approvals

We consider baseline T1- and T2-weighted MR images (resolution 1 mm) from an iNPH patient collected in a previous clinical study. This patient then also underwent a (0.5 ml, 1 mmol/ml) intrathecal injection of gadobutrol, and follow-up MR images were taken at several time points post injection. LookLocker images were also obtained with the T1-weighted MR images. The clinical study was approved by the Regional Committee for Medical and Health Research Ethics (REK) of Health Region South-East, Norway (2015/96), the Institutional Review Board of Oslo University Hospital (2015/1868), the National Medicines Agency (15/04932-7), and conducted in accordance with the ethical standards of the Declaration of Helsinki of 1975 (and as revised in 1983). All study participants were included after written and oral informed consent.

### 2.2 In-vivo imaging concentration estimates

The baseline MR images were post-processed using FreeSurfer v6.0^38^ to obtain a segmentation of the brain. To define a choroid plexus (CP) completely enclosed by the lateral ventricles, a CP domain was manually marked in the images. Next, the left and right pial membrane, white matter interface, cerebellum, ventricles and aqueduct were represented via triangulated surfaces. The segmentation of the SAS was performed by thresholding a registered T2-weighted image, and any clusters not connected to the FreeSurfer segmentation were removed. Subsequently, a surface bounding the SAS was constructed, and expanded by 1 mm in the surface normal direction to ensure that the SAS was represented as a continuous compartment between the pia and dura around the whole brain. The CSF volume before and after expansion were 457 and 602 mL, respectively. The spinal cord was not segmented, and was represented as CSF for simplicity. The parenchymal volume was 1266 mL. Both the CSF and parenchymal volumes are slightly above average values in iNPH patients^31^.

The generated surfaces were further post-processed using SVMTK^39^, and finally used to generate a volumetric mesh Ω of the parenchyma Ω_*P*_ and surrounding CSF-spaces Ω_*F*_ combined (Figure 1). We label the boundary separating Ω_*P*_ and Ω_*F*_ by *∂*Ω_*P*_. The choroid plexus Ω_CP_ ⊂ Ω_*F*_ is located within the lateral ventricles and we denote its surface (in contact with the CSF) by *∂*Ω_*CP*_. The outer boundary of the SAS is split into three parts: *∂*Ω_*S*_, *∂*Ω_FM_, and *∂*Ω_out_, representing the arachnoid membrane, foramen magnum and a chosen efflux route, respectively. We consider and define three different regions Ω_out_ for efflux of CSF: locally across the *parasagittal dura* (Figure 2**A**), locally across the *cribriform plate* (Figure 2**B**), or into the meningeal *lymphatics* distributed over the outer (arachnoid) boundary (Figure 2**C**). Finally, to simulate retro-grade net aquaductal flow, we consider flow into the choroid plexus (Figure 2**E**) from the foramen magnum (Figure 2**D**).

**Figure 2.**
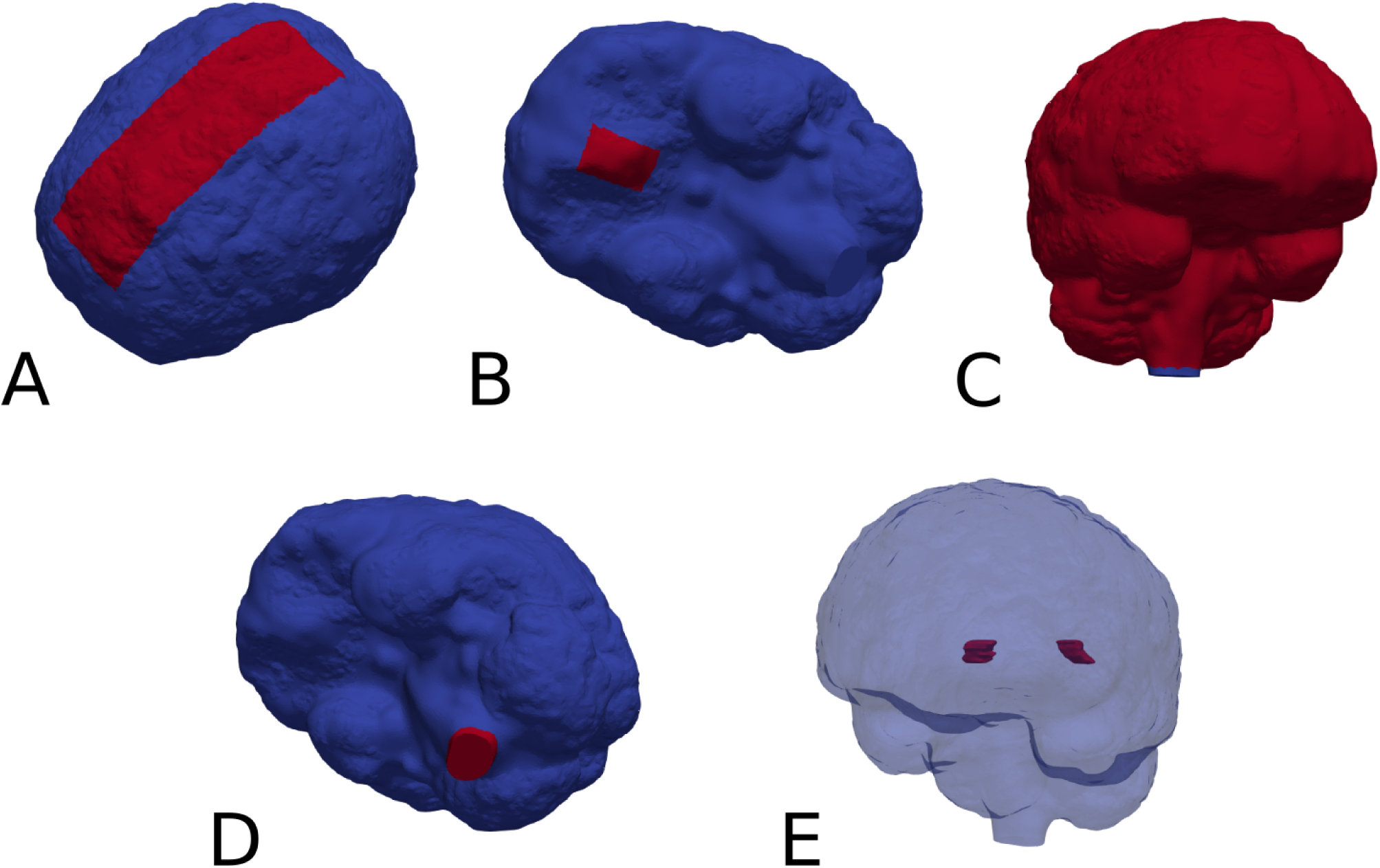
Red markers highlight important subregions and boundaries in the computational domain: the **(A)** parasagittal dura, **(B)** cribriform plate, **(C)** meningeal lymphatics, **(D)** foramen magnum and **(E)** choroid plexus.

### 2.3 Flow in the CSF spaces

We model the flow of CSF in Ω_*F*_ by the incompressible Stokes equations: find the CSF velocity field *u* and pressure *p* such that

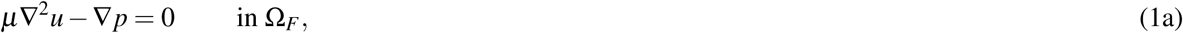

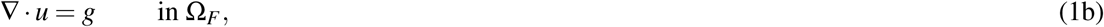

where *g* is a given source of fluid. With the low Reynolds numbers (0.001) reported for flow in PVS^9, 40^, we find steady Stokes flow to be a reasonable assumption for the present study. To represent CSF production in the choroid plexus, we let *g* be a given positive constant in Ω_*CP*_ and zero elsewhere in Ω_*F*_. Specifically, by default, we set *g* such that approximately 0.5L of CSF is produced every 24 hours. We also consider a scenario with increased CSF-production. In humans, the CSF production has been reported to increase during sleep^41, 42^, while high CSF turnover through lymphatics has been reported in awake mice^7^. We set the parenchymal CSF/brain interstitial fluid (ISF) velocity to be zero (in Ω_*P*_).

We set the CSF velocity at the outer boundary (representing the arachnoid membrane) to be zero, except at specific efflux/absorption sites *∂*Ω_out_ to be further specified. At these, we set a traction condition:

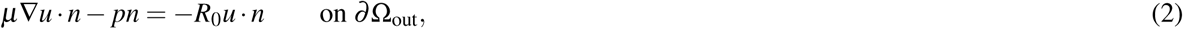

where *R*_0_ *≥* 0 represents an efflux resistance acting to moderate CSF outflow in these regions, and *n* denotes the outward pointing boundary normal. The fluid source, in combination with the zero or low resistance efflux routes, induces a flow of CSF from the CP through the ventricular system, through the SAS and out across either the parasagittal dura, cribriform plate, or meningeal lymphatics.

We also consider a reversed flow scenario, in which *g* is set negative with a value corresponding to a sink of 0.5 L/day, a zero traction condition is imposed at the foramen magnum *∂*Ω_FM_, and zero velocity (no slip) is imposed on the remainder of the boundary.

### 2.4 Molecular transport in the CSF and parenchyma

We also model molecular transport within the CSF-spaces and parenchyma resulting from an influx of gadobutrol at the foramen magnum (resulting e.g. from an intrathecal injection). We model transport of a concentration *c* in the entire domain Ω via the diffusion-convection equation:

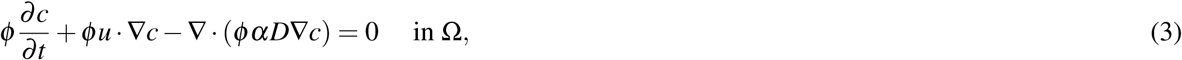

where *u* is a convective velocity field, *D* denotes an apparent diffusion coefficient, and *α* is a dispersion factor. We set the apparent diffusion coefficients *D*_*F*_ = 3.8 10^−4^ mm^2^/s in Ω_*F*_ and 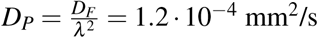 in Ω_*P*_^28^. Here, *λ* ≈ 1.78, represents the tortuosity. To represent enhanced diffusion in the CSF due to pulsatile effects, mixing or other forms of dispersion^23, 29, 43^, we have introduced the dispersion factor *α*, and consider a range of *α* ∈ {1, 10, 100, 1000} in Ω_*F*_. In Ω_*P*_ we set *α* = 1. *ϕ* accounts for the porosity of the extracellular space which occupies 20 % of the parenchyma^44^, and we thus set *ϕ*_*P*_ = 0.2 and *ϕ*_*F*_ = 1. We consider either *u* = 0 and *α* = 1 (diffusion-only scenarios) or let *u* be given by solutions of the CSF flow equations (1) in combination with all *α*.

To represent an influx of gadobutrol at the foramen magnum, we set

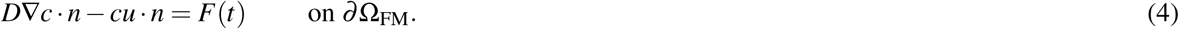

Based on tracer enhancement as reported by Eide et al.^6^, *F*(*t*) is modeled as a linearly decreasing function until *T*_0_ *≈* 2.24 hours (8064 s) and zero thereafter i.e.

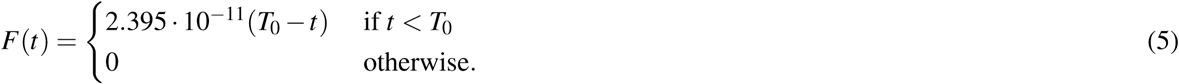

The solute influx *F*(*t*) (given in mmol/(s mm^2^) is chosen such that the total amount of gadobutrol injected is approximately 0.5 mmol. At the efflux sites *∂*Ω_out_, we let the solute be absorbed via the relation

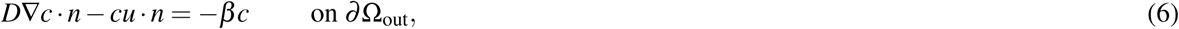

where *β* is a given membrane permeability. The case *β* = 0 corresponds to no absorption, *β* = ∞ corresponds to free movement of solutes across the boundary, while 0 < *β* < ∞ represents a diffusive resistance to molecular outflow. On the remainder of the boundary, we do not allow for solute efflux, by setting *D*∇*c n cu n* = 0. Moreover, we let the initial concentration be *c*(*x*, 0) = 0. Note that to model transport associated with the reversed flow scenario, we let *∂*Ω_CP_ take the role of *∂*Ω_out_.

At the interface between Ω_*F*_ and Ω_*P*_ we conserve mass (enforce conservation of molecules) by setting *ϕD*_*P*_∇*c*_*P*_*·n* = *D*_*F*_ ∇*c*_*F*_*·n*. Here, *D*_*P*_ and *D*_*F*_ denote *D* restricted to Ω_*P*_ and Ω_*F*_, respectively, *n* is the normal vector on the interface, pointing from Ω_*P*_ to Ω_*F*_ and *ϕ* denotes the ECS porosity.

### 2.5 Overview of models

CSF and solutes may have several simultaneous and possibly partially independent outflow routes^2^. We here consider six different flow and transport models separately (Table 1), each with different dispersion factors. This design allows us to systematically examine different pathways and evaluate whether each or combinations could describe in-vivo observations of Gadobutrol transport. Model I and II describe flow induced by CSF production in the CP, and CSF efflux across the parasagittal dura and cribriform plate, respectively. For these models, we assume free molecular efflux at the absorption sites. Model III is a variant of Model I with a finite molecular efflux permeability at the parasagittal dura absorption site. Model IV reflects a different efflux pathway with CSF production in the CP, CSF efflux int the meningeal lymphatics, and a finite molecular efflux permeability. Model V represents a reversed flow scenario with absorption of CSF in the CP region (and CSF influx at the foramen magnum). Finally, Model VI represents a variant of Model II with increased CSF production.

**Table 1.** Overview of computational models. Production and absorption site refers to production site for CSF and efflux/absorption site of CSF and the solute concentration, respectively. *R*_0_ is a CSF efflux resistance parameter cf. (2), while *β* represents a diffusive resistance to molecular efflux cf. (6). The values for *R*_0_ and *β* were estimated by numerical experimentation.

| Model | Production site | Absorption site | $R_0$ (Pa/(mm s)) | $\beta$ (mm<sup>2</sup>/s) | Production (L/day) |
| --- | --- | --- | --- | --- | --- |
| I | Choroid plexus | Parasagittal dura | 0 | ∞ | 0.5 |
| II | Choroid plexus | Cribriform plate | 0 | ∞ | 0.5 |
| III | Choroid plexus | Parasagittal dura | 0 | $10^{-4}$ | 0.5 |
| IV | Choroid plexus | meningeal lymphatics | $10^{-5}$ | $10^{-4}$ | 0.5 |
| V | Foramen magnum | Choroid plexus | 0 | ∞ | 0.5 |
| VI | Choroid plexus | Cribriform plate | 0 | ∞ | 1.0 |

### 2.6 Numerical methods, simulation software and verification

The Stokes equations are solved using a finite element method with Taylor-Hood (continuous piecewise quadratic and continuous piecewise linear) elements for the velocity and pressure. The diffusion-convection equation with boundary conditions is solved numerically using the finite element method with continuous linear finite elements for the concentration in space, and the backward Euler method in time; all using the FEniCS finite element software^45, 46^. The brain mesh has 6 691 432 cells and 1 088 640 vertices. The degrees of freedom for the diffusion equation is equal to the number of vertices. For the Taylor-Hood case the number of degrees of freedom is 27 858 018. Moreover, the largest cell size is 2.4 mm and the smallest is 0.07 mm. The largest cells are in the middle of white matter where there are no stokes flow or sharp gradients.

A time resolution study was performed to ensure that our simulation results were independent of the choice of time step (Supplementary Figure S1). Including testing and validation, a total of *≈* 30,000 CPU hours was used to run the simulations on big memory nodes. All simulations were run on the high-performance computing infrastructure Sigma2 – the National Infrastructure for High Performance Computing and Data Storage in Norway.

### 2.7 Concentration estimates from in-vivo MRI

We extract contrast agent concentration estimates from the MR images post injection for comparison with computational predictions. The contrast agent shortens the T1 times as:

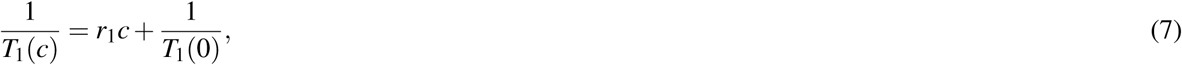

where c denotes the concentration of the contrast agent, *r*_1_ is known as the T1 relaxivity of the agent, and *T*_1_(*c*) and *T*_1_(0) denote the T1 time with and without concentration, respectively. The T1 times can be computed using a T1 mapping^47^, such as the LookLocker sequence. Through a preliminary phantom study, the relaxivity constant for this LookLocker protocol was found to be 6.5 L mmol^−1^ s^−1^. The median T1 time over the parenchyma was used in (7) to estimate the concentration in the parenchyma. The CSF concentration was estimated by manually creating a region of interest (ROI) in the CSF, and using the average T1 time over the ROI with (7). Finally, to transform the concentration in the parenchyma to be that of the the extracellular space, the concentrations was multiplied by five.

### 2.8 Quantities of interest

The total amount of solute in a given region Ω_*i*_ (*i* = *F, P*) at time *t* was computed as 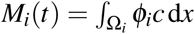. The total amount within the intracranial compartment *M*(*t*) is then the sum *M*(*t*) = *M*_*P*_(*t*) + *M*_*F*_ (*t*). The average concentrations per region over time were computed as

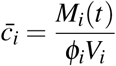

where *V*_*i*_ refers to the volume of the respective region. To compare parenchymal influx between models, we compute the peak average concentration in the parenchyma and the time to reach this peak. We also compute the relative clearance of tracers after *T*_1_ = 3 days as 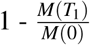

## 3 Results

All models induce non-trivial CSF flows through the ventricular system and subarachnoid space.

### 3.1 Different outflow routes induce different CSF flow patterns and velocities

Models I–IV all reach maximal SAS velocities of 8.9 mm/s in the thinnest part of the aqueduct (Figure 3**A**-**C**). Despite their differences in efflux pathways, all of these models predict higher CSF flow velocities in the anterior regions of the SAS compared to posterior regions. Model II displays the highest velocities in the SAS, reaching 50 *μ*m/s. Model I (and III) reaches peak CSF velocities of 40 *μ*m/s. In model IV, CSF flow occurs mainly in the lower regions of the SAS as CSF can exit the SAS along the entire boundary. Peak velocities in the SAS for model IV reach 20 *μ*m/s. In models where CSF was allowed to exit through outflow routes other than the parasagittal dura (models II and IV), CSF velocity magnitudes were relatively small (< 4 *μ*m/s) in the SAS near the upper convexities of the brain.

**Figure 3.**
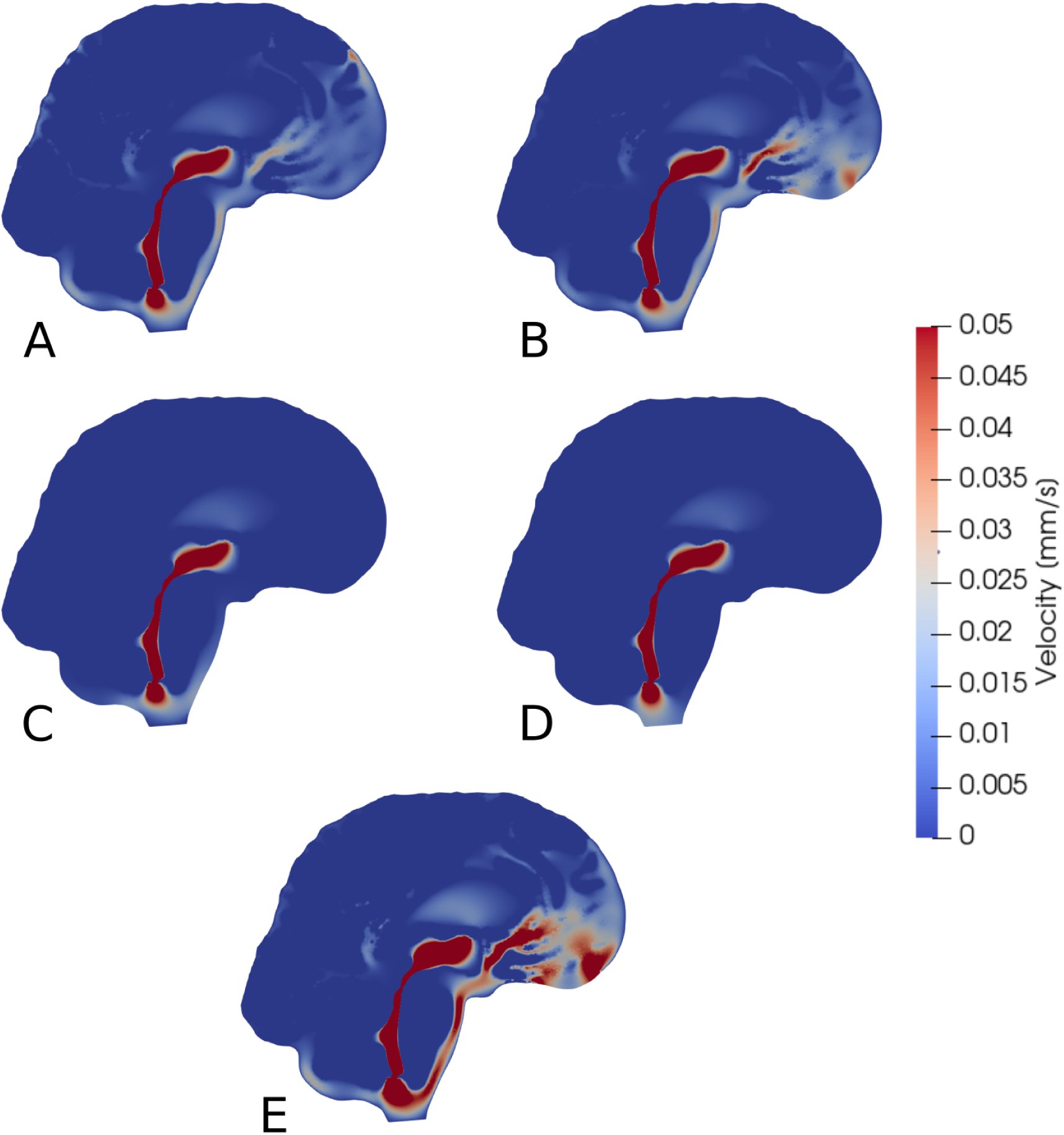
Sagittal views (cut through the center of the aqueduct) of CSF velocity magnitudes induced by steady CSF production in the choroid plexus combined with different CSF efflux pathway models, or a reversed flow scenario. Subfigures show velocity fields resulting from CSF efflux through **(A)** the parasagittal dura, **(B)** the cribriform plate, **(C)** the meningeal lymphatics **(D)** production in the foramen magnum and absorption in the choroid plexus, or **(E)** the cribriform plate with double production. The color map is capped at 0.05 mm/s for visualization purposes.

### 3.2 Reversed CSF flow pathways

Model V predicts that, under its assumptions, CSF will predominantly flow from the foramen magnum directly to the CP, limiting CSF flow in other parts of the SAS (Figure 3**D**). thus the flow direction is reversed compared to models I–IV. In the foramen magnum, CSF velocity magnitudes reach 20 *μ*m/s, while the velocity in the aqueduct remain at 8.9 mm/s. In the upper regions of the SAS, not directly associated with the 3rd ventricle, CSF velocities were typically lower than 0.1 *μ*m/s.

### 3.3 Increased CSF production increase CSF velocities

Doubling the CSF production (model VI versus model II) results in a doubling of the CSF velocity field by linearity. Therefore, we observe velocities of approx. 100 *μ*m/s in the CSF space (Figure 3**E**) and a velocity in the aqueduct of 17.8 mm/s for model VI.

### 3.4 Diffusion alone yields excessively slow clearance from intracranial compartments

When driven purely by diffusion (without convection or dispersion enhancements), the tracer spreads radially from the foramen magnum and distributes evenly throughout the brain. Distribution is slightly faster in the CSF than in the parenchyma, as the free diffusion coefficient in CSF is larger. However, this effect is not very noticeable. For Models I–III the relative one year clearance is only 32.8 %, 17.6 % and 29.9 %. Model IV displays faster clearance, clearing 92.5 % over one week, but with a late peak parenchyma concentration occurring after 79 hours.

### 3.5 Tracer distribution patterns induced by CSF circulation and dispersion

Including the CSF circulation-induced flow as a convective velocity substantially speeds up the clearance rates, both from the SAS and parenchyma.

Tracer distribution is shown for all models after 6 and 24 hours and *α* = 10 in Figure 4, revealing substantial inter-model variations. For Model I, the tracer is mainly confined to the SAS and moves upwards towards the parasagittal dura showing a clear preference traveling along the SAS in the right hemisphere (data not shown). As there is no molecular molecular resistance to outflow on the parasagittal dura in Model I, tracer is instantly transported out when moving into this efflux route. In regions where the tracer concentration in the SAS is high, the tracer also enters the brain due to the large concentration gradient between the SAS and the brain (4A, B Model I). After one week, a some tracer are still found within the brain, slowly diffusing back towards the pial surface for clearance via convection in the SAS (data not shown). Models I and III (with outflow via the parasagittal dura) are the only models where tracer reaches the upper convexities of the brain, resulting in a brain wide distribution of tracers. In Model III, where a diffusive molecular resistance is added at the parasagittal dura, tracer accumulates near the outflux region (4B, Model III).

**Figure 4.**
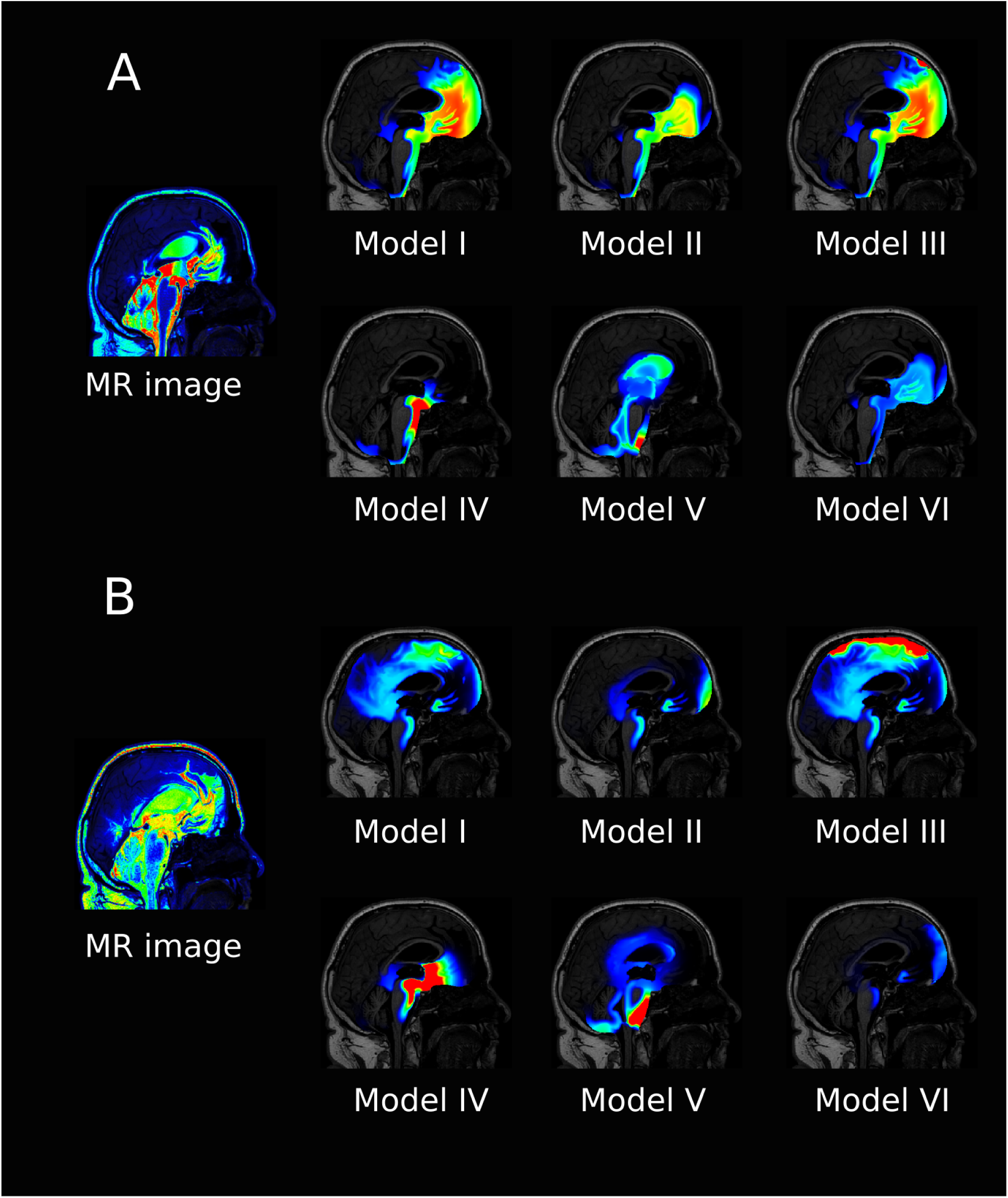
The figure shows a sagittal view of all the models at 6 hours (**A**) and 24 hours (**B**) after intrathecal injection of gadobutrol for *α* = 10. For the simulation data, the colorscale shown is 0.1–5 mmol/L in **A** and 0.1-1 mmol/L in **B**. For comparison the T1 contrast enhanced image for the patient at the same times are included. The MR images are scaled separately for picture legibility.

Model V is the only model where tracer reaches the ventricular system, while Model IV has localized accumulation of tracers around the brain stem. Model VI, with increased CSF production, show generally lower concentration of tracers, and some accumulation near the outflux route at the cribriform plate.

The average concentration over time for all models, and *α* = 10, is compared in Figure 5, both for the ISF and CSF. The figure also contains in vivo concentration estimates in both spaces. We observe that a combination of the different outflow routes, i.e. Model I and V, gives a comparable result to that of the MR images. Model I and III both display higher concentrations than the data in both the CSF and parenchyma/ISF (Figure 5). Model II, IV and V, on the other hand, yield the comparable or lower concentrations.

**Figure 5.**
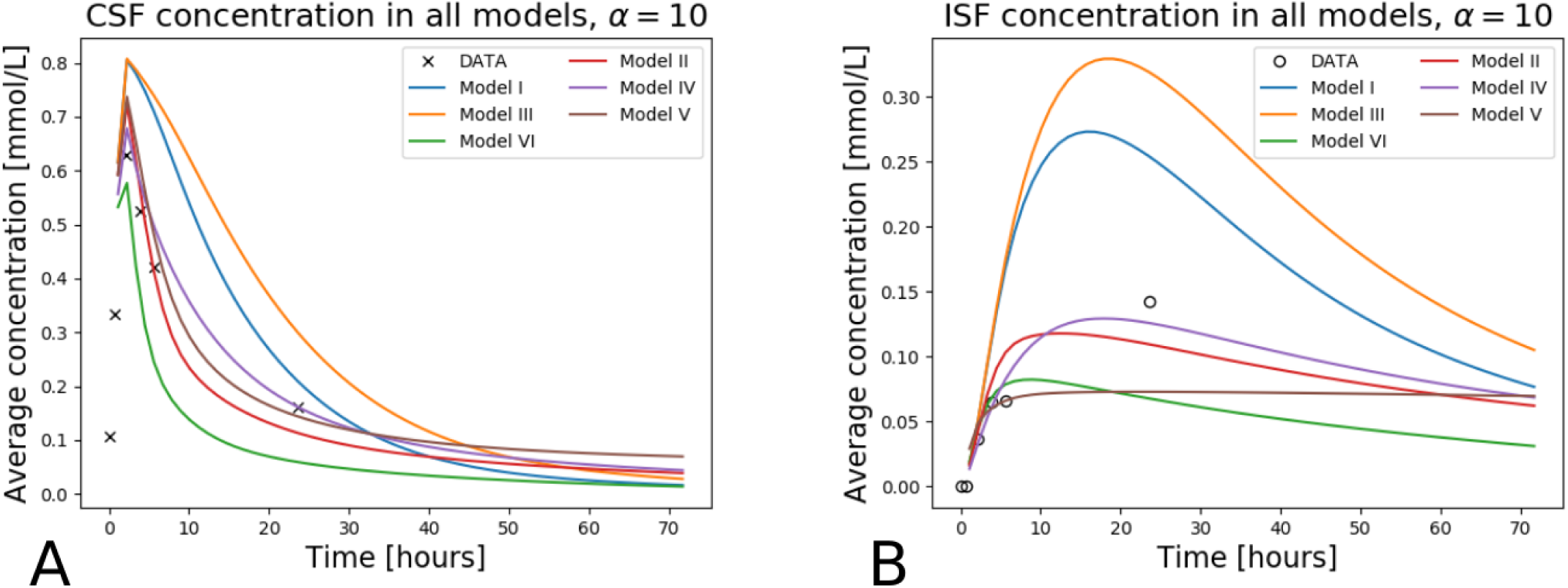
The figure shows concentration in the CSF **(A)** and the ISF **(A)** for all models over one week. The tracer concentration data from T1 MR images for this patient is also shown.

### 3.6 Clearance rates induced by CSF circulation and dispersion

Model I and II both display high three-day clearance rates for all dispersion factors (Figure 6**A**-**B**). Specifically, the three day clearance rates are between 94.1 and 97.7 % for Model I and between 88.9 and 94.9 % for Model II (Table 2). The tracer concentration is initially higher in the SAS allowing for diffusive influx to the brain. At later time points, the SAS has been cleared, mainly via convective flow, and the tracer partly remains inside the parenchyma, delaying the total clearance of tracers from the intracranial compartment. Model I has slightly higher peak average parenchyma concentration values than Model II, reaching 0.30 and 0.24 mmol/L, respectively. The time to peak in the parenchyma occurred after 7.8 – 19.0 hours for Model I and 10.1 – 16.8 hours or Model II.

**Figure 6.**
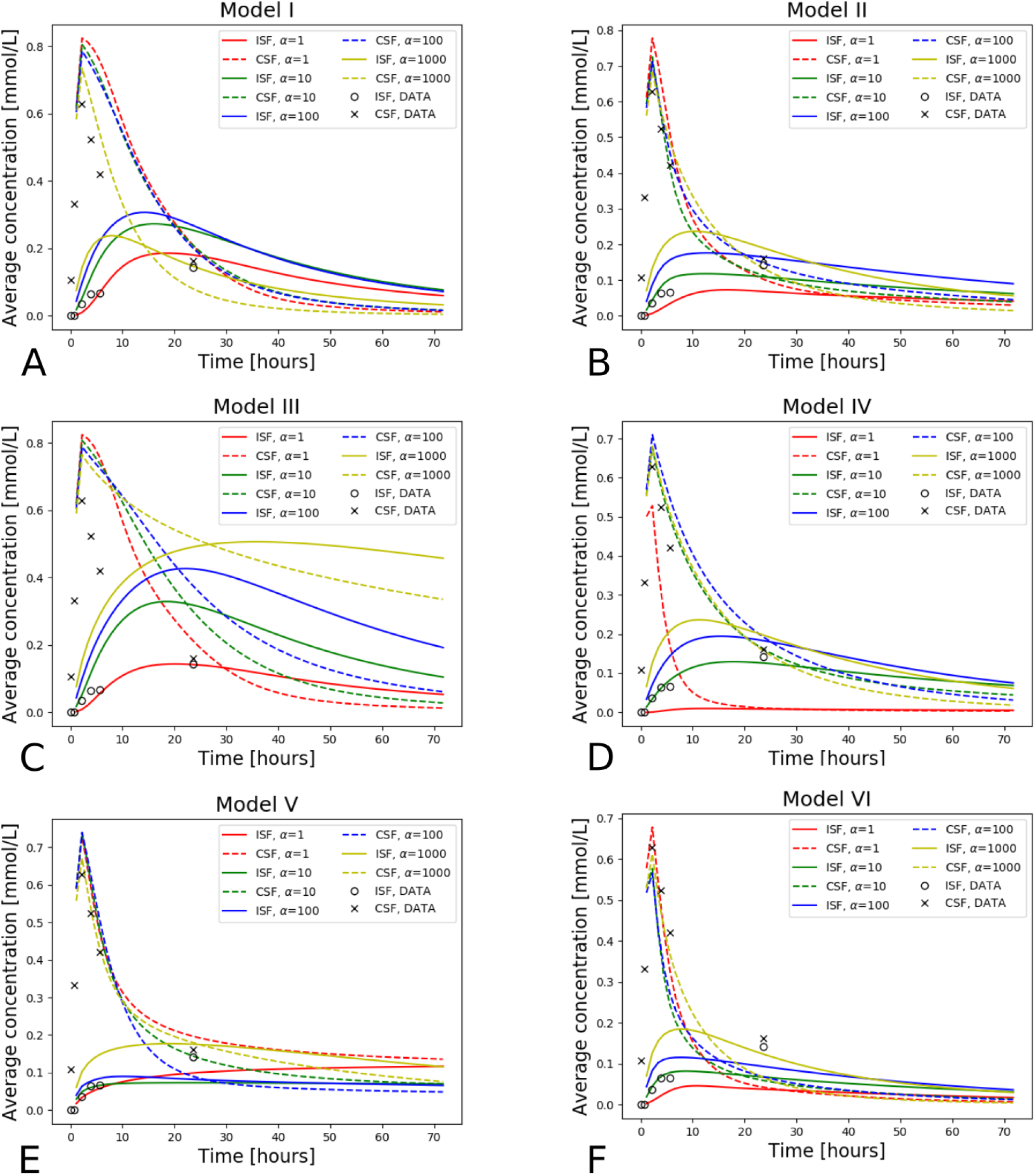
Average concentration in the parenchyma (par.) and CSF over a period of 72 hours. Models I-VI is used with dispersion values *α* = 1, 10, 100, 1000. Also plotted is the concentration data taken from T1-weighted images of this specific patient as a ground truth. The tracer injection (present from 0 to 2.24 hours) is seen as a sharp increase in CSF concentration at early time points. When the injection is no longer present, the total amount of tracers within the intracranial compartment starts decreasing. The tracer concentration data from T1 contrast enhanced images for the patient is also included. **(A)** Model I, **(B)** Model II, **(C)** Model III, **(D)** Model IV, **(E)** Model V and **(F)** Model VI.

**Table 2.**
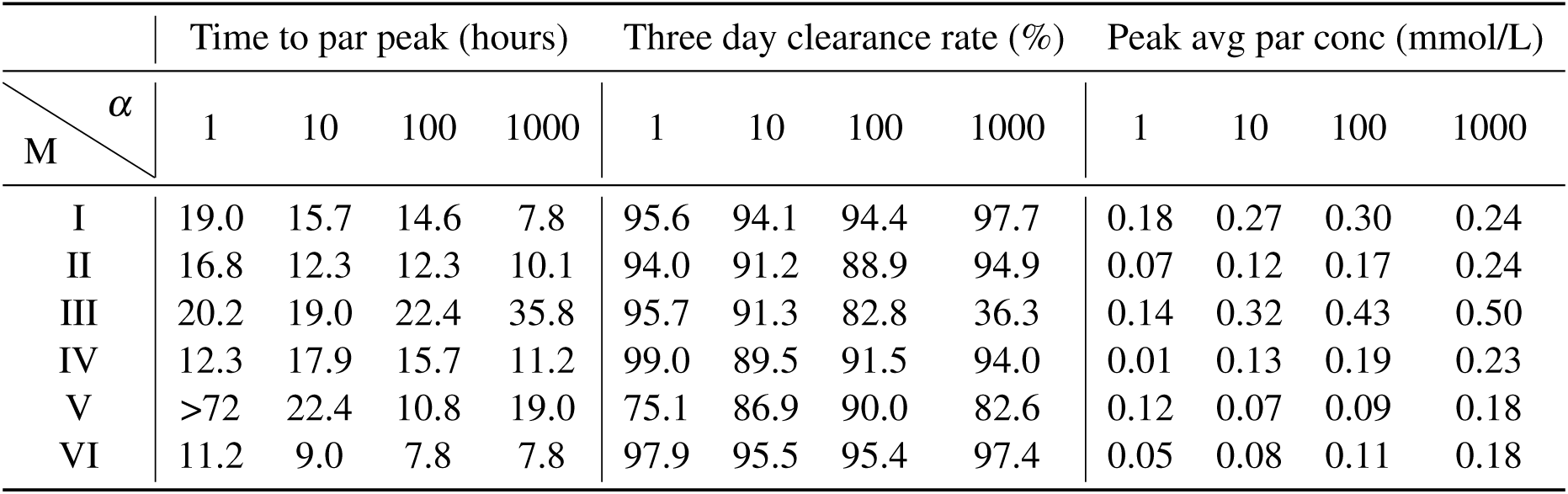
The table shows time to peak concentration in the parenchyma (left), total mass clearance in the intracranial compartment after 72 hours (middle) and peak average concentration values in the parenchyma (right) for the case of gadobutrol transport. Values are shown for all models and *α* values. M: Model, *α*: Dispersion factor, par: parenchyma, conc: concentration, avg: average

| M \ $\alpha$ | Time to par peak (hours) | | | | Three day clearance rate (%) | | | | Peak avg par conc (mmol/L) | | | |
| --- | --- | --- | --- | --- | --- | --- | --- | --- | --- | --- | --- | --- |
|  | 1 | 10 | 100 | 1000 | 1 | 10 | 100 | 1000 | 1 | 10 | 100 | 1000 |
| I | 19.0 | 15.7 | 14.6 | 7.8 | 95.6 | 94.1 | 94.4 | 97.7 | 0.18 | 0.27 | 0.30 | 0.24 |
| II | 16.8 | 12.3 | 12.3 | 10.1 | 94.0 | 91.2 | 88.9 | 94.9 | 0.07 | 0.12 | 0.17 | 0.24 |
| III | 20.2 | 19.0 | 22.4 | 35.8 | 95.7 | 91.3 | 82.8 | 36.3 | 0.14 | 0.32 | 0.43 | 0.50 |
| IV | 12.3 | 17.9 | 15.7 | 11.2 | 99.0 | 89.5 | 91.5 | 94.0 | 0.01 | 0.13 | 0.19 | 0.23 |
| V | >72 | 22.4 | 10.8 | 19.0 | 75.1 | 86.9 | 90.0 | 82.6 | 0.12 | 0.07 | 0.09 | 0.18 |
| VI | 11.2 | 9.0 | 7.8 | 7.8 | 97.9 | 95.5 | 95.4 | 97.4 | 0.05 | 0.08 | 0.11 | 0.18 |

For the models including a molecular resistance to outflow at the outflow site (i.e. Model III and IV), the three day clearance rate is comparable to Models I and II, except for the case when *α* = 1000 in model III (Table 2). The highest three day clearance is obtained with *α* = 1 for both Model III and IV (95.7 and 99.0 % clearance, respectively). The lowest three day clearance is obtained with *α* = 1000 for model III (36.3 % clearance) and *α* = 10 for model IV (89.5 % clearance, Table 2). Model III reaches a peak parenchyma concentration of 0.50 mmol/L, while Model IV have a lower peak of 0.23 mmol/L. The time to peak exceeds 19.0 hours for all dispersion factors in Model III, which is much later than the other models. Model IV on the other hand peaks between 11.2 and 17.9 hours.

### 3.7 Clearance of gadobutrol with reverse pathways

Model V (with reversal of CSF flow in the aqueduct) results in low parenchymal enrichment compared to Models I–IV (Table 2, Figure 6E). The three day clearance rate is between 75.1 and 90.0 % depending on *α* and the peak average concentration is 0.18 mmol/L in the parenchyma (Table 2). The time to peak concentration in the parenchyma is long for *α* = 1, occurring later than after 1 week, but for larger dispersion factors, the peak occurs between 10.8 and 22.4 hours.

### 3.8 Increased CSF production results in rapid clearance

Model VI, with double the CSF production of the other models, displayed rapid clearance from the CSF (Figure 6**F**). The rapid turnover of CSF limited the influx and facilitated clearance also within the parenchyma. The three day clearance rate for all dispersion factors ranged between 95.4 – 97.9 % (Table 2). The peak average parenchyma concentration occurs early, between 7.8 and 11.2 hours and reaches at most 0.15 mmol/L, when *α* = 1000.

## 4 Discussion

In this paper we have simulated molecular transport by diffusion and convection for six different models investigating distribution of gadobutrol molecules entering the intracranial compartment via the foramen magnum. The different models represent different outflow routes, and CSF flow patterns vary considerably between models. The effect of outflow route and dispersion factor modify the distribution and clearance patterns in a non-linear and unpredictable manner. Outflow through either the parasagittal dura, the cribriform plate or through meningeal lymphatics, typically cleared 80–99% of injected tracers over a time period of three days. These three models however, display very different spatial distribution of tracers. In Models I and III tracer distributes more or less throughout the frontal cortex, while when outflow occurs through meningeal lymphatics, tracers are mainly located at the brain stem at the base of the brain.

With a daily production of 0.5 and 1.0 L/day in our models, the velocity reaches 10 and 17.8 mm/s in the aqueduct. Pulsatile aqueductal flow velocities of several cm/s have been measured experimentally^48–50^ in the range 1-10 cm/s. Average velocities max velocities were reported around 5 cm/s, corresponding to a total volume flux of 0.3 mL per cycle, of which 0.01 mL was net^4, 32^. As the net flow is around 1/30 of the total flux, the corresponding net max velocity can then be estimated as 5/30 cm/s, somewhat below the velocities estimated here. Further, in iNPH patients it has been reported Phase-contrast MR has reported retro-grade net flow in the aqueduct^4^. Model V is motivated by retro-grade aqueductal flow and we see that this model is distinct from the other models in that there is significant ventricular enrichment, as often seen in iNPH^15^.

On the pial surface of the brain, we observe velocities up to 20 – 50 *μ*m/s in models I-IV, and up to 100 *μ*m/s in model VI. These velocities align relatively well with experimentally observed bulk flow velocities of around 20 *μ*m/s observed in mice^8, 9^. Thus, CSF flow observed in these studies, may very well be a result of CSF production and absorption, driven by small static pressure differences. It should be noted that mice have approximately 3x faster CSF turnover compared to humans^36^. Given otherwise similar CSF dynamics between the species, one would thus expect CSF production to cause higher velocities at the surface of the mouse brain compared to a human. Comparing model II and model VI, the increased CSF efflux to the cribriform plate limits tracer influx to the brain, in line with the hypothesis of Ma et al.^7^. We observe that models with a short distance between injection and absorption site (models II, IV and V) limits the influx of tracers to the parenchyma. In general, tracers will enter the brain if present on the surface over a long time period. For a given tracer, the amount of tracers entering the brain will thus be affected by both the CSF velocity and the distance from the injection site to the absorption site.

Gadobutrol injections has been studied in human subjects in several papers. Eide, Ringstad and colleagues have reported MR intensity increase for a large number of subjects^6, 15, 22, 35^, while Watts et al^51^ quantified gadobutrol concentrations over time in a single patient. These studies show an initial sharp increase in tracer concentration in the SAS, typically reaching a peak at around 2-6 hours. In the parenchyma, peak values occur between 10 and 24 hours, depending on the region of interest. Gray matter regions, closer to the pial surface typically peak at around 10 hours, while for specific white matter regions, peak values may occur closer to 24-40 hours post-injection^15, 51^. In all our models, peak CSF concentration occurs at the time when the gadobutrol influx at the foramen magnum is turned off, i.e. after approximately 2 hours. More interestingly, the ISF concentration peaks later, and the time to peak is between 10 and 20 hours in 15 out of 24 models tested. ISF concentration is reported to decay relatively slow, with an approximate concentration at 48 hours at half its peak value^22^. Furthermore, the peak concentration of gadobutrol has been measured as 0.5 mmol/L in the CSF and around 0.12 mmol/L in the ISF^51^, in line with both the estimates of observed concentration the iNPH patient and the results from our models. Both Model II (outflow through the cribriform plate) and Model IV (outflow via meningeal lymphatics) match all these criteria well when the dispersion in the SAS was modeled by *α* = 10. Model I (outflow through the parasagittal dura) replicate experimentally observed ISF concentration without additional dispersion in the SAS, but clearance from the SAS is delayed in this model compared to experimental data. With a molecular resistance to outflow on the parasagittal dura (Model III), simulations reproduce accumulation of tracers in this region, but clearance kinetics are slower than expected. With a doubling of CSF production (Model VI) the kinetics of ISF and CSF clearance is faster than expected for all dispersion factors tested. In the model with reversed flow in the aqueduct (Model V), we qualitatively reproduce the tracer enhancement in the aqueduct as seen in iNPH patients^6^. However, rapid flow through the aqueduct and into the choroid plexus prevents the expected brain-wide enhancement of tracer^15^, and in Model V tracers are confined to the foramen magnum or in the vicinity of the lateral ventricles. Combined, these results suggest that a combination of production and efflux sites may be needed to reproduce the observed tracer distribution^6, 15, 22^.

The role of different outflow routes from the SAS have been debated and challenged over years. In particular, the traditional view of outflow predominately through arachnoid granulations has been criticized recently^2^. Our Model I is conceptually similar to outflow through arachnoid granulations with CSF draining close to the dural sinus. The results from our simulations can not exclude any of the proposed major outflow routes, as all of them resemble experimental data in at least some measure. A specific weighting between inflow and outflow routes may potentially be sufficient to explain differences between groups (e.g. iNPH vs control), or differences between individuals. The results do show unequivocally that CSF flow and clearance is a major player in CNS clearance. Convective flow in the SAS speeds up intracranial clearance from years to hours and days, an enormous effect compared to the effect of bulk flow of around 1 *μ*m/s within the ECS^27^. Furthermore, changes in the dispersion factor (increased diffusion due to mixing) only in the SAS changed both peak values and clearance rates within the brain ECS.

In terms of limitations, we only performed the simulations on a single patient. To create one patient specific mesh with high mesh quality that includes all anatomical regions of interest was time consuming, and increasing the amount of subjects was not the scope of this study. Similarly, as the initial mesh already consisted of nearly 7M cells, a mesh resolution study was not performed. Based on previous simulation studies^27^, using similar number of cells, we believe the mesh is sufficiently resolved. To resolve all regions of the SAS, the SAS was expanded by 1 mm. This modification increases the volume of which fluid flows, and thus slightly reduce velocities we find in the SAS. The total CSF volume was increased by around 33%, we thus assume that our reported SAS flow velocities of 20–50 *μ*m/s are lower estimates. In the SAS, we assumed that the dispersion factor were similar in all subregions. In reality, dispersion would be expected to be enhanced close to larger arteries^35^ and in regions where pulsatile CSF flow is substantial (e.g. near the foramen magnum). Furthermore, we did not include ISF velocities in the foramen magnum. There is very little knowledge exactly on how the velocity fields are directed^27^, especially without a priori knowledge of the location of blood vessels. In addition, the purpose of this study was to assess the effect of SAS convection, independent of potential bulk flow within the brain. Finally, we should note that we assumed that all injected gadobutrol reached the foramen magnum, while around 33 % of CSF has been proposed to be drained along the spinal canal^52^. The latter point may explain the fact that most of the reasonable models tested (Model I, II and IV) all generally display a slight overestimation of the SAS peak concentration in our models compared to the data.

In conclusion we have demonstrated that convection in the SAS yield rapid clearance both from the SAS and the ISF, even when pure diffusive transport were assumed in the ECS. Convective fluid flow in the SAS has the potential to speed up clearance from years (as would be the case for purely diffusive transport) to days. As none of the models tested were able to reproduce the observed data perfectly (both qualitatively and quantitatively), a combination of the different outflow routes seem most plausible, and their relative weight may differ between groups^6^.

## Author Contributions

M.H., V.V., L.M.V., P.K.E, G.R., M.E.R. and K.A.M. conceived the simulations, L.M.V. and M.H. segmented and meshed MR images, M.H. conducted the simulations, V.V. M.H. and L.M.V did the analysis of the results and M.H. made the figures. All authors discussed the simulations and results. M.H., V.V, L.M.V., M.E.R. and K.A.M. wrote the first draft. All authors revised and approved the final manuscript.

