## Supplementary material for "CSF circulation and dispersion yield rapid clearance from intracranial compartments"

### 1 SUPPLEMENTARY DATA

#### 1.1 Time resolution convergence

In Figure S1 we present the first 10 hours of simulation results from Model I using a time step of $\Delta t = 4000$, $\Delta t = 2000$ and $\Delta t = 1000$. The results shown in the Results section in our paper was obtained using $\Delta t = 4032$. The error always stayed below 3.7 % in the SAS and 6.5 % in the ISF relative to the peak concentration observed in model I.

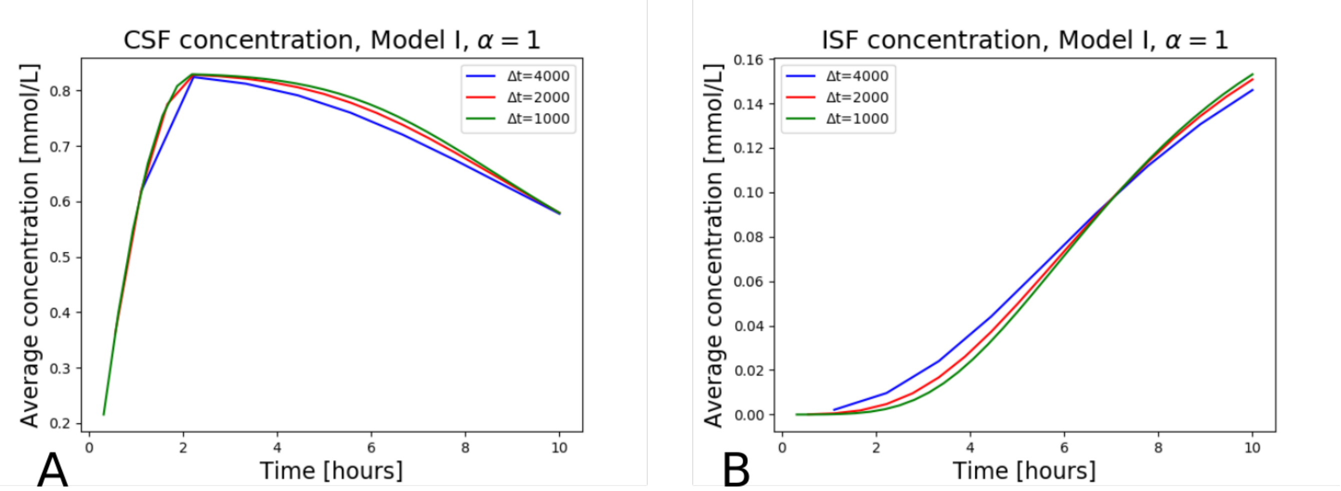


**Figure S1.** The figure shows the CSF and ISF average concentration in Model I with $\alpha = 1$ for the first 10 hours. Plotted are the results for three different time steps of increasing resolution.
